# Dementia is Associated with a Syndrome of Global Neuropsychiatric Disturbance

**DOI:** 10.1101/813485

**Authors:** Donald R. Royall, Raymond F. Palmer, the Alzheimer’s Disease Neuroimaging Initiative

## Abstract

**Objective:** Global factors have been identified in measures of cognitive performance (i.e., Spearman’s *g*) and psychopathology (i.e., “General Psychopathology”, “*p*”). Dementia is also strongly determined by the latent phenotype “δ”, derived from *g*. We wondered if the Behavior and Psychological Symptoms of Dementia (BPSD) might arise from an association between δ and *p*.

**Methods:** δ and *p* were constructed by confirmatory factor analyses in data from the Alzheimer’s Disease Neuroimaging Initiative (ADNI). δ and orthogonal factors representing “domain-specific” variance in memory (MEM) and executive function (EF) were regressed onto p and orthogonal factors representing “domain-specific” variance in positive (+) and negative (-) symptoms rated by the Neuropsychiatric Inventory Nursing Home Questionnaire (NPI-Q) by multiple regression in a structural equation model (SAM) framework.

**Results:** Model fit was excellent (CFI = 0.98, RMSEA = 0.03). δ was strongly associated with *p*, (+) and (-) and strongly associated with *p* (r = −0.57, p<0.001). All three associations were inverse (adverse). Independently of δ, MEM was uniquely associated with (+), while ECF was associated with (-). Both associations were moderately strong. ECF was also weakly associated with *p*.

**Conclusions:** Dementia severity (δ) derived from general intelligence (*g*) is specifically associated with general psychopathology (*p*). This is *p*’s first demonstration in an elderly sample and the first to distinguish the global behavioral and psychological symptoms *specific to dementia* (BPSSD) from behavioral disturbances arising by way of non-dementing, albeit likely disease-specific, processes affecting domain-specific cognitive and behavioral constructs. Our findings call into question the utility of proposed regional interventions in BPSSD, and point to the need to explore global interventions against dementia-specific behavioral features.

## Introduction

The so-called Behavioral and Psychological Symptoms of Dementia (BPSD) are a major source of comorbidity and caregiver burden (1). BPSD increase exposure to psychopharmacological agents and their associated adverse drug reactions (ADR) (2). They are a major factor in caregiver stress (3-4), increase the risk of institutionalization (5) and inflate the costs associated with dementia care (3).

Regardless, surprisingly little is known about the biological substrates of BPSD, their natural history or risk factors. It is widely assumed that BPSD arise from the regional neuropathology(ies) specific to the disease(s) in which they develop. Some investigators have attempted to use to BPSD to develop etiologically precise BPSD signatures. However, in young adults, a global “general psychpathology” factor “*p*” has been proposed to influence all psychopathological behavioral domains (6-7) and is demonstrable in a wide range of conditions (8).

*p* has never been addressed in the context of dementia and BPSD. However, it recapitulates Spearman’s well-accepted general intelligence factor “*g*” (9). Our recent work strongly implicates *g* as the essential cognitive determinant of dementia severity (10). Only via *g* can cognitive performance be meaningfully related to functional status. We have demonstrated this via theory-driven bi-factor confirmatory analyses (CFA) in a Structural Equation Model (SEM) framework (11-12). The resulting latent variable “δ” (for “dementia”) is strongly associated with the Clinical Dementia Rating Scale (CDR) “Sum of boxes” (CDR-SB) (13), cross-sectionally (14), longitudinally (15-16), and across diagnoses (16).

δ is highly accurate in distinguishing demented cases (16), but it is entirely “agnostic” as to dementia’s etiology (17). If *p* can be shown to be associated specifically with δ then it would directly implicate dementia *itself* as the transdiagnostic determinant of global psychopathology, link *p* to intelligence, and constrain *p*’s biology to that of *g* and δ.

Conversely, “domain-specific” variance in cognitive performance (i.e., “memory”, “frontal lobe functions”, etc.) is “orthogonal” (unrelated) to *g* (11-12). It therefore risks to be unrelated to both δ and IADL, and thus to be independent of dementia *per se*. In this study, we test the fit of *p* to BPSD reported participants in the Texas Alzheimer’s Research and Care Consortium (TARCC) and relate cognitive performance, including δ and orthogonal domain-specific factors, to prospective variance in *p* and orthogonal domain-specific behavioral clusters.

## Methods

### Subjects

Subjects included n = 3385 participants in the Texas Alzheimer’s Research and Care Consortium (TARCC). The Consortium’s methods have been described in detail elsewhere (18). Briefly, the TARCC cohort is a convenience sample of well-characterized cases of Alzheimer’s disease (AD) (n = 1240), “Mild Cognitive Impairment “(MCI) (n = 688), and normal controls (NC) (n = 1384). Each TARCC participant undergoes a standardized annual examination that includes a medical evaluation, neuropsychological testing, and clinical interview. Diagnosis of AD is based on National Institute for Neurological Communicative Disorders and Stroke-Alzheimer’s Disease and Related Disorders Association (NINCDS-ADRDA) criteria. All TARCC evaluations and psychometrics are provided in English or in Spanish according to the subject’s preference. Institutional Review Board approval was obtained at each site and written informed consent was obtained for all participants.

### Statistical Analyses

This analysis was performed using Analysis of Moment Structures (AMOS) software (19). The maximum likelihood estimator was chosen. Co-variances between the residuals were estimated if they were significant and improved fit.

#### Neuropsychiatric Indicators

*The Neuropsychatric Inventory (NPIQ) (20):* Behavioral and neuropsychiatric disturbances were measured at Wave 2 by the informant-rated NPI-Q. Multiple Spanish translations are available. We used the Spanish version generated by the Spanish Translation and Adaptation Work Group (STAWG) from the National Alzheimer’s Coordinating Center (NACC) Uniform Data Set (UDS) (21). This scale has been well-validated among Mexican American (MA) respondents (22).

The NPI-Q encompasses 12 behavioral features that are commonly exhibited in the context of neuropsychiatric illness. For this study, a modified NPIQ was administered by an experienced research assistant to knowledgeable informants (caregivers), who were asked to rate the presence and the severity of each BPSD, i.e., agitation/aggression, dysphoria/depression, irritability/lability, apathy/indifference, anxiety, disinhibition, aberrant motor behavior, delusions, hallucinations, euphoria/elation, nighttime behavioral disturbances, and appetite/eating disturbances.

Informants were first asked to endorse the presence or absence of each BPSD over the prior four weeks, using a single screening question. If the informant endorsed the behavior, a severity score was assigned on a three-point Likert scale (i.e., 1-mild, 2-moderate, 3-severe). If the informant did not endorse a behavior, its “severity” was rated “zero”. Thus, in our modified adaptation, each NPI-rated BPSD ranged from zero (not endorsed) to 3 “severe”. The total NPI-Q severity score represents the sum of the 12 individual modified symptom severity scores and ranges from 0 to 36 (NPI-QTOT).

#### Covariates

All observed measures in the structural models were adjusted for baseline age, body mass index (BMI), education, ethnicity, gender, Mini-Mental Status Exam (MMSE) score (23), **HCY, and HgbA1c.**

##### Age

Self-reported age was confirmed by birthdate and coded continuously.

##### Body Mass Index (BMI)

BMI was estimated as the ratio of subject height to weight (REF).

##### Education

Education was coded continuously as years of formal education.

##### Ethnicity

Ethnicity was determined by self-report and coded dichotomously as “Hispanic” = 1 and “non-Hispanic White” (NHW) = 0.

##### Gender

Gender was coded dichotomously with “female” = 1.

The Clinical Dementia Rating Scale sum of boxes (CDR-SB) (24): The Clinical Dementia Rating Scale “sum of boxes (CDR-SB) (13) The CDR is used to evaluate dementia severity. The rating assesses the patient’s cognitive ability to function in six domains – memory, orientation, judgment and problem solving, community affairs, home and hobbies and personal care. Information is collected during an interview with the patient’s caregiver. Optimal CDR-SB ranges corresponding to global CDR scores are 0.5 - 4.0 for a global score of 0.5, 4.5 - 9.0 for a global score of 1.0, 9.5 - 15.5 for a global score of 2.0, and 16.0 - 18.0 for a global score of 3.0.

The Geriatric Depression Rating Scale (GDS) (25): GDS scores range from zero-30. Higher scores are worse. A cut-point of 9-10 best discriminates clinically depressed from non-depressed elderly.

The Mini-Mental Status Examination (MMSE) (23): The MMSE is a well-known and widely used test for cognitive impairment screening. Scores range from 0 to 30. Scores less than 24 reflect cognitive impairment.

#### Analysis Sequence

The NPI-Q items’ severity ratings were submitted to a CFA in SEM. First, a latent variable “*p*” was derived from all 12 BPSD. Orthogonal latent factors rating “positive” (+) and negative (-) domain-specific symptom clusters were constructed from *p*’s residuals. This renders (+) and (-) “orthogonal” (unrelated) to *p* because their respective indicators are already adjusted for that global construct. The result is a “bifactor” CFA model defining *p* and two orthogonal domain-specific factors, (+) and (-) (**Figure 1**).

**Figure 1:**
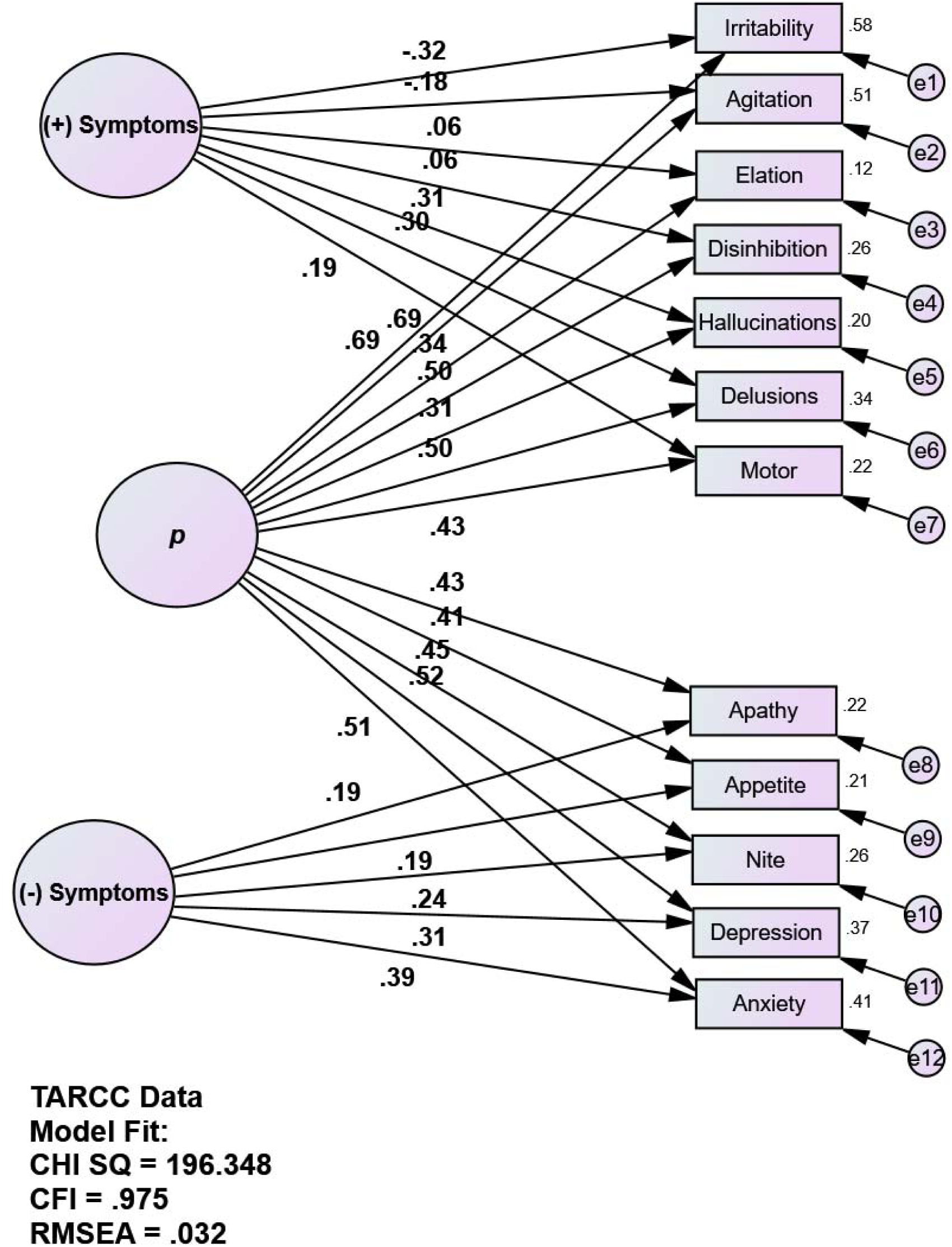
Bifactor CFA of BPSD. Index; BPSD = Behavior and Psychological Symptoms of Dementia; CFA = Confirmatory Factor Analysis; GDS = Geriatric Depression Scale; HCY = serum homocysteine; HgbA1c = serum hemoglobin A1c; RMSEA = Root Mean Square Error of Association. *All observed variables except APOE are adjusted for age, education, ethnicity, gender, GDS, HCY, and HgbA1c (paths not shown for clarity). Those covariates are densely intercorrelated.

The bifactor model’s fit was compared to several alternatives, beginning with the single factor *p*. In a second alternative, (+) and (-) were used as *p*’s indicators, resulting in a hierarchical model. Finally, all three models containing *p* were compared to a previously described four factor model which lacked this construct. It was developed by an exploratory factor analysis of NPI-Q data (26). All four alternative models were constructed from NPI-Q data collected at wave 2.

The best fitting moldel was associated with cognitive performance data collected at wave 1 in TARCC (baseline). Now, cognitive performance data were submitted to a four factor CFA as previously described (12). This results in the “dDx” δ homolog, and three orthogonal factors, i.e., g’ (dDx’s residual in Spearmans’s *g*) and two domain-specific cognitive factors rating memory (MEM) and executive function (EF). In TARCC, the dDx homolog has been reported to have a high area under the receiver operating characteristic curve (AUC /ROC) for Alzheimer Disease’s (AD)’s discrimination from normal controls (NC) (c = 0.98) and to be strongly associated with dementia severity as measured by the CDR-SB (r = 0.88).

dDx, MEM and EF at wave 1 were regressed onto wave 2 *p*, (+) and (-) by multivariate regression in a structural SEM framework. The result is a multivariate regression model of the baseline cognitive factors as predictors of prospective global and domain-specific BPSD (**Figure 2**).

**Figure 2:**
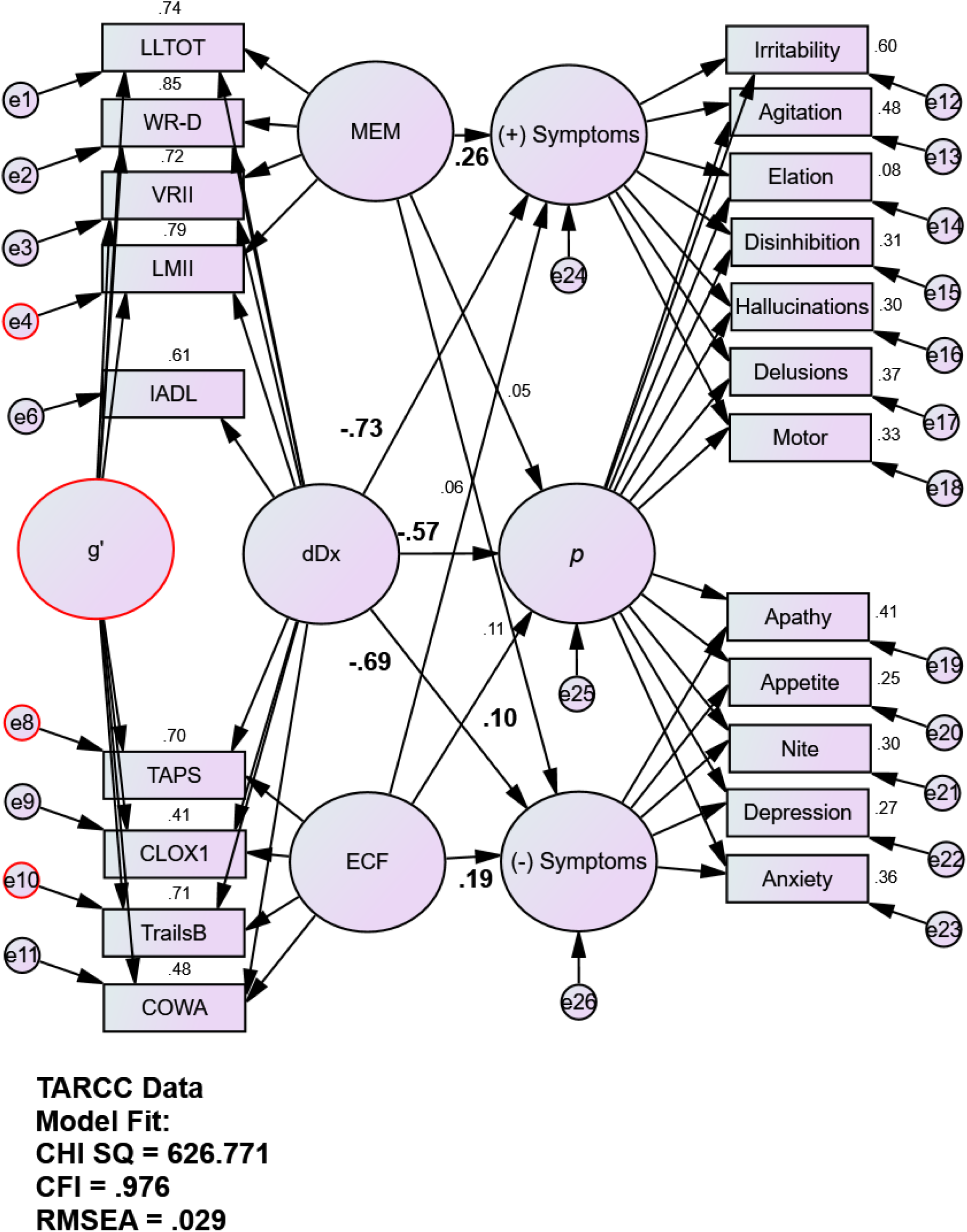
dDx is Strongly Associated with *p*. Index; GDS = Geriatric Depression Scale; HCY = serum homocysteine; HgbA1c = serum hemoglobin A1c; RMSEA = Root Mean Square Error of Association. *All observed variables except APOE are adjusted for age, education, ethnicity, gender, GDS, HCY, and HgbA1c (paths not shown for clarity). Those covariates are densely intercorrelated.

#### Missing data

We used the newest instance of TARCC’s dataset (circa 2016). The entire dataset was employed. Clinical diagnoses were available on 3385 subjects, 2861 of whom had complete data for δ’s cognitive indicators and covariates. Modern Missing Data Methods were automatically applied by the AMOS software (27). AMOS employs Full information Maximum Likelihood (FIML) (28).

#### Fit indices

Fit was assessed using four common test statistics: chi-square, the ratio of the chisquare to the degrees of freedom in the model (CMIN /DF), the comparative fit index (CFI), and the root mean square error of approximation (RMSEA). A non-significant chisquare signifies that the data are consistent with the model (29). However, in large samples, this metric conflicts with other fit indices (insensitive to sample size) show that the model fits the data very well. A CMIN/DF ratio < 5.0 suggests an adequate fit to the data (30).The CFI statistic compares the specified model with a null model (31). CFI values range from 0 to 1.0. Values below 0.95 suggest model misspecification. Values approaching 1.0 indicate adequate to excellent fit. An RMSEA of 0.05 or less indicates a close fit to the data, with models below 0.05 considered “good” fit, and up to 0.08 as “acceptable”(32). All fit statistics should be simultaneously considered when assessing the adequacy of the models to the data.

## Results

The demographic characteristics of TARCC’s sample are presented in Table 1.

**Table 1:** Descriptive Statistics.

| Variable | N | Mean (SD) |
| --- | --- | --- |
| Age (observed) | 3381 | 70.88 (9.48) |
| CDR (Sum of Boxes) | 3306 | 2.42 (3.35) |
| EDUC (observed) | 3381 | 13.24 (4.25) |
| Ethnicity (1 = MA, n = 1189) | 3381 | 0.36 (0.47) |
| GDS<sub>30</sub> (observed) | 3005 | 5.60 (5.25) |
| Gender (♂ = 1, n = 1281) | 3312 | 0.39 (0.49) |
| IADL (Summed) | 3381 | 10.48 (4.52) |
| MMSE | 3311 | 25.52 (4.76) |
| Complete Cases | 2861 |  |

CDR = Clinical Dementia Rating scale; COWA = Controlled Oral Word Association Test; DIS = Digit Span Test; GDS = Geriatric Depression Scale; IADL = Instrumental Activities of Daily Living; MMSE = Mini-mental State Exam; SD = standard deviation; WMS LM II = Weschler Memory Scale: Delayed Logical Memory; WMS VR I = Weschler Memory Scale: Immediate Visual Reproduction.

**Table 2:** Model Fit.

| Alternative Models | CHISQ | df | CFI | RSMEA | ACC |
| --- | --- | --- | --- | --- | --- |
| 1 *p* only | 551.55 | 54 | 0.92 | 0.05 | 623.55 |
| 2 Hierarchical | 463.32 | 52 | 0.93 | 0.05 | 539.32 |
| 3 Conventional (no *p*) [REF] | 2677.81 | 53 | 0.58 | 0.12 | 2751.81 |
| 4 Orthogonal bifactor<br>[*p*, (+), (-) (**Figure 1**)] | 196.35 | 42 | 0.98 | 0.03 | 292.35 |

Base model fit was excellent (CFI = 0.98, RMSEA = 0.02). All behaviors loaded significantly on *p*. The *ad hoc* (+) and (-) factors were significantly indicated by their respective BPSD. The base model had improved fit over all alternatives.

The base model’s three BPSD factors, *p*, (+) and (-), were regressed onto baseline psychometric performance. Model fit was again excellent (CFI = 0.98, RMSEA = 0.03). δ was strongly associated with p, (+) and (-). All three associations were inverse (adverse). Independently of δ, MEM was uniquely associated with (+), while ECF was associated with (-). Both associations were moderately strong. ECF was also weakly associated with *p*.

## Discussion

This analysis demonstrates a global BPSD factor’s relatively good fit to observed BPSD and *p*’s statistically strong association with dementia severity as measured by δ. Additionally, MEM was associated specifically with (+), while ECF was associated with (-), and with dementia.

Since δ and *g* are both “indifferent to their indicators”, they manifest in a wide range of cognitive performance measures. We have demonstrated δ’s psychometric stability across batteries, and even down to the item-set of an individual measure (33). The spectrum of CNS-variables impacted by δ and *g* may extend beyond cognition, into sensory or even motor function. Spearman himself constructed a *g* homolog from sensory discrimination tasks alone (9).

Those findings appear to constrain δ’s (*and therefore dementia’s*) biomarkers to those of “intelligence” (Caveat: Since *g* is manifest in *all* cognitive performance measures and possibly in sensory and motor domains, “intelligence” in this context cannot be construed as meaning “cleverness”, “rationality” or “wisdom”. Spearman’s use of the word is limited to a unique mathematical association, demonstrable by factor analysis). Dementia’s true biomarkers must be similarly positioned to impact *every cognitive performance measure*. Regional pathologies are disadvantaged in this regard.

The present findings suggest a similar constraint may relate to BPSD. Global psychopathology (i.e., *p*) may relate to the globally distributed processes that impact *g* through δ. Domain-specific behavioral constructs, orthogonal to *p*, seem more likely have regional determinants (34-35). This is consistent with our findings that MEM is related to (+) but not (-), while ECF is related to (-) and not (+). MEM might be plausibly associated with regional mesio-temporal pathology while ECF can be plausibly associated with regional frontal circuit pathology. Replication of this analysis by the dT2A homolog in ADNI (14) might allow for confirmation of such hypotheses against structural and functional neuroimaging.

The possibility of a dementia-specific dysbehavioral syndrome was alluded to by Hughlings Jackson in his 1884 description of the “dissolution” of the central nervous system (CNS) (36). He argued for a hierarchical loss of neurological integrity that would disrupt higher abilities and thereby “release” lower centers from control.

> “I submit that disease only produces negative mental symptoms answering to the dissolution, and that all elaborate positive mental symptoms (illusions, hallucinations, delusions, and extravagant conduct) are the outcome of activity of nervous elements *untouched by any pathological process*” (emphasis added). -Hughlings Jackson, 1884

Jackson further described “uniform” and “local” dissolutions. “In uniform dissolution the whole nervous system is under the same conditions or evil influence, the evolution of the whole nervous system is comparatively evenly reversed.” He recognized that local pathologies arise from distinct etiologies and that “Different kinds of insanity are different local dissolutions of the highest centres.” This suggests that the behaviors arising from global CNS disturbances might be a transdiagnostic property of dementing illness. The regional disruptions unique to each dementing illness might engender diagnostically-specific behavioral features, but those would not present transdiagnostically.

In this scenario, a global dysbehavioral syndrome would emerge in demented states regardless of their etiology(ies) and in response to transdiagnostic pathology, not disease-specific changes. Support for transdiagnostic dementia-specific behavioral features is provided by the stereotyped natural history of BPSD at specific stages of dementia’s evolution (37), the transdiagnostic presentation of multiple BPSD (38-39) and high penetrance of BPSD with increasing dementia severity (40). Support for a global vs. “local” BPSD dichotomy can also be intuited in recent work on the bifactor nosology of psychopathology in children and adults. Factor analyses of behavioral psychopathology suggest independent effects of both global and domain-specific behavioral factors on observed psychopathology (8).

Our model also suggests the potential for a novel *rational approach* to the treatment of BPSD. BPSD arising from regional pathology(ies) and manifesting as domain-specific behavioral issues may respond to “regional” pharmaco-biological interventions. BPSD manifesting as *p*-related behavioral issues may require “global” interventions and /or interventions directed at δ and its biomarkers.

Examples of regional interventions might include many traditional monoaminergic approaches. Monoaminergic networks arise in the brainstem and project to regionally precise targets in the neocortex and associated subcortical structures (41). They do not project globally throughout the brain. Examples of monoaminergic interventions for BPSD might include atypical antipsychotics and Serotonin Selective Reuptake Inhibitor (SSRI) antidepressants. Even-the pro cholinergic cholinesterase inhibitors are reported to have efficacy against certain BPSD (42). More precisely localized regional interventions would include regional lobotomy, deep-brain stimulation of specific structures (e.g., the amygdala), and /or regional transcranial magnetic stimulation (rTMS). All have been advocated for BPSB and related psychobehaviors in the literature (43).

In contrast, γ-aminobutyric acid (GABA) and glutamate are ubiquitously distributed, as are the effects of hypoglycemia, seizure disorders, whole brain radiation, and possibly blast-related traumatic brain injuries (TBI). Alcohol (EtOH) and benzodiazepines (BNZ) may have GABA-mediated adverse effects on behavior and cognition manifesting as changes in *g*, δ and *p*. The protean impacts of those insults beyond cognition and behavior, *including also balance, sensory and motor performance* may betray effects on *g*, δ and *p*. Thus, the BPSD-related to *g*, δ and *p* may respond better to global interventions, e.g., mood stabilizing anticonvulsants, lithium, and or electroconvulsive therapies (ECT). Alternatively, *g*, δ and *p* might be adversely impacted by interventions of this class, explaining the insalubrious reputation of BNZ and EtOH in cognitively impaired persons.

In summary, we have specifically associated general psychopathology (*p*) with a latent dementia severity metric (δ) derived from general intelligence (*g*). This is *p*’s first demonstration in an elderly sample and the first to distinguish the global behavioral and psychological symptoms *specific to dementia* (BPSSD) from behavioral disturbances arising by way of non-dementing, albeit disease-specific processes affecting domain-specific cognitive and behavioral constructs. Our findings call into question the utility of proposed regional interventions in BPSSD, and point to the need to explore global interventions against dementia-specific behavioral features.

## Acknowledgments

This work was supported by the Julia and Vann Buren Parr endowment for the study of Alzheimer’s Disease. The funders had no role in study design, data collection and analysis, decision to publish, or preparation of the manuscript.

Some data used in preparation of this article were obtained from the ADNI database (adni.loni.usc.edu). Data collection and sharing for this project was funded by the Alzheimer’s Disease Neuroimaging Initiative (ADNI) (National Institutes of Health Grant U01 AG024904) and DOD ADNI (Department of Defense award number W81XWH-12-2-0012). ADNI is funded by the National Institute on Aging, the National Institute of Biomedical Imaging and Bioengineering, and through generous contributions from the following: AbbVie, Alzheimer’s Association; Alzheimer’s Drug Discovery Foundation; Araclon Biotech; BioClinica, Inc.; Biogen; Bristol-Myers Squibb Company; CereSpir, Inc.; Cogstate; Eisai Inc.; Elan Pharmaceuticals, Inc.; Eli Lilly and Company; EuroImmun; F. Hoffmann-La Roche Ltd and its affiliated company Genentech, Inc.; Fujirebio; GE Healthcare; IXICO Ltd.; Janssen Alzheimer Immunotherapy Research & Development, LLC.; Johnson & Johnson Pharmaceutical Research & Development LLC.; Lumosity; Lundbeck; Merck & Co., Inc.; Meso Scale Diagnostics, LLC.; NeuroRx Research; Neurotrack Technologies; Novartis Pharmaceuticals Corporation; Pfizer Inc.; Piramal Imaging; Servier; Takeda Pharmaceutical Company; and Transition Therapeutics. The Canadian Institutes of Health Research is providing funds to support ADNI clinical sites in Canada. Private sector contributions are facilitated by the Foundation for the National Institutes of Health (www.fnih.org). The grantee organization is the Northern California Institute for Research and Education, and the study is coordinated by the Alzheimer’s Therapeutic Research Institute at the University of Southern California. ADNI data are disseminated by the Laboratory for Neuro Imaging at the University of Southern California.

This analysis has been presented as an abstract at the Gerontological Society of America’s 71st Annual Scientific Meeting in Austin, Texas on November 16, 2019.

